# InsectMorphoAI: A Deep Learning-Based Software for Automated Estimation of Insect Length, Volume, and Biomass

**DOI:** 10.1101/2025.05.22.655251

**Authors:** Hossein Shirali, Aleida Ascenzi, Lorenz Wührl, Nils Beyer, Noemi di Lorenzo, Emanuele Vaccarella, Nathalie Klug, Rudolf Meier, Pierfilippo Cerretti, Christian Pylatiuk

## Abstract

We present InsectMorphoAI, an open-source, user-friendly software package that automates the measurement of insects from 2D images. The software addresses the need for high-throughput, non-invasive alternatives to laborious and often destructive manual measurement methods. InsectMorphoAI provides two analysis modes: a rapid, general-purpose method using oriented bounding boxes for linear length estimation across diverse taxa, and a high-precision, taxon-specific instance segmentation method for detailed curvilinear length, volume, and biomass estimation. We demonstrate the software’s accuracy, showing that the volume estimates from the segmentation module are strongly correlated with dry weight (R = 0.907), and the general length module achieves a mean absolute error corresponding to ~2.3% of the average specimen length. InsectMorphoAI is distributed with a graphical user interface and is freely available, with straightforward installation via a Docker container or a native Python environment. By streamlining data acquisition, InsectMorphoAI facilitates the integration of detailed trait data into large-scale ecological research, from biodiversity monitoring to functional trait analysis.

## 1. Introduction

Accurate measurement of insect traits, such as length, volume, and biomass, is essential for studying ecosystem health, species distribution, and ecological dynamics. Such traits provide important insights into a wide array of biological processes, from food webs, energy flow, and how insect populations respond to environmental changes. This also includes the decline of insect populations across ecosystems whose quantification required efficient morphometric tools to monitor changes and inform conservation strategies (Hallmann et al., 2017; Sánchez-Bayo & Wyckhuys, 2019). Accurate biomass data for model species are equally in demand because body size governs predator–prey encounter rates, handling times, and food-web structure (Brose, 2010; Miller-ter Kuile et al., 2022) and is central to estimating reproductive output and progeny size (Fox & Czesak, 2000). Pollination studies rely on precise body mass information to quantify foraging range, pollen-load capacity, and the fit between pollinators and flowers (Bartomeus et al., 2016; Greenleaf et al., 2007). Species-level measurements are also essential for tracking population trends (Lister & Garcia, 2018) and for interpreting allometric scaling in functional trait analyses (Cariveau et al., 2016; Kendall et al., 2019). Yet traditional weighing of individual insects is destructive, slow, and often impossible for tiny or rare taxa. However, obtaining morphometric data remains a significant challenge. A notable example is the study of bulk samples collected using methods such as Malaise traps (Karlsson et al., 2020). Traditional methods often involve letting wet samples drip-dry to a consistent state or direct weighing after oven-drying; both approaches are invasive, labour-intensive, and time-consuming.

To address these challenges, the field is rapidly moving towards integrated pipelines that combine robotics, imaging, high-throughput barcoding, and machine learning to tackle large-scale insect analysis (Wägele et al., 2022). This includes automated specimen handling (Wührl et al., 2022), size-sorting (Ascenzi et al., 2025), megabarcoding (Meier et al., 2025), high-resolution imaging (Hereld et al., 2017; Klug et al., 2024), and AI-driven classification (Caruso et al., 2025; Shirali et al., 2024). The “reverse workflow” in molecular studies, where specimens are barcoded first and then morphologically validated (Hartop et al., 2024; Srivathsan et al., 2021), further emphasizes the need for efficient, non-destructive morphometric tools. While systems like BIODISCOVER (Ärje et al., 2020), DiversityScanner (Wührl et al., 2022), and other deep learning approaches (Schneider et al., 2022) have pioneered automated biomass estimation, a gap remains for a flexible, specimen-level tool that offers both broad applicability and taxon-specific precision.

Here, we introduce InsectMorphoAI, an open-source software package designed to fill this gap, offering key advantages over existing systems by providing (1) a rapid and high-precision method; (2) a focus on user-friendliness through an intuitive graphical interface and simple Docker deployment; and (3) the ability to output detailed, specimen-level metrics such as curvilinear body length and segmented part volumes. This paper introduces the application, describes its dual-mode functionality, and demonstrates its accuracy, showing how it facilitates the integration of trait data into high-throughput biodiversity monitoring workflows.

## 2. Software Description

### 2.1. Overview and Availability

InsectMorphoAI is a Python-based application featuring a web-based graphical user interface (GUI) built with the Streamlit framework (Streamlit Inc., n.d.), designed for ease of use by researchers without a computational background. The software is open-source (MIT License) and publicly available on GitLab. To ensure broad accessibility and reproducibility, we provide two installation methods. The primary method uses a Docker container, which encapsulates all dependencies and allows for a straightforward, one-command setup on any major operating system (Windows, macOS, Linux). For users who need to process large batches of images or require more control, we provide detailed instructions for a native Python installation, as well as a command-line interface. The repository includes a comprehensive README.md file with step-by-step installation guides, usage examples, and a small dataset for testing the setup. While the graphical interface is designed to run on standard personal computers, processing large datasets with the command-line interface is most efficient on systems with substantial RAM (≥16 GB) and a dedicated NVIDIA GPU.

### 2.2. The User Workflow

The interface is designed for flexibility and ease of use, organized into a compact side panel with expandable sections and a main panel for results (Figure 1).

**Figure 1.**
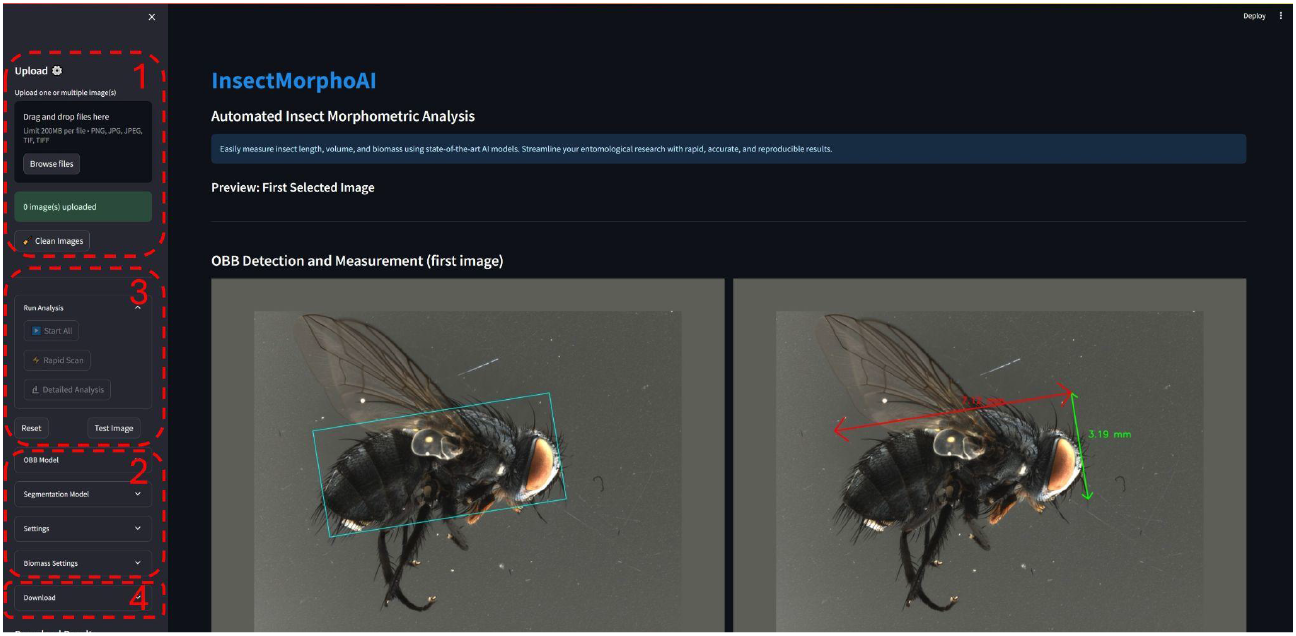
The InsectMorphoAI graphical user interface (GUI). Key areas: (1) Image upload and management; (2) Expandable “Settings” for camera, lens, and biomass parameters; (3) “Run Analysis” buttons; and (4) “Download Options” for exporting results.

#### (1) Image Upload & Management

The user begins by uploading one or more images. Utility buttons enable the quick removal of all uploaded files (“Clean Images”) or the loading of a pre-packaged test image to instantly demonstrate the full analysis workflow. The underlying models were trained on high-quality images of ethanol-preserved specimens captured using the Entomoscope (Wührl et al., 2024), which have clean, uniform backgrounds. The models can process specimens from various camera views, not limited to a strict lateral perspective, and users can expect the best performance with images that share these characteristics.

#### (2) Parameter Configuration

In the “Settings” panel, the user can configure all analysis parameters. For image calibration, the software includes a list of predefined camera and lens profiles from the Entomoscope imaging system, each with a stored pixel-to-millimeter conversion factor. A “Custom Setup” option also allows users to define their own camera, lens, and conversion factor. In the “Biomass Estimation” section, users can enable or disable this calculation and edit the default taxon-specific parameters to match their own data.

#### (3) Analysis Selection

The “Run Analysis” section provides three distinct options: a “Rapid Scan” button for general length estimation, a “Detailed Analysis” button for taxon-specific morphometrics, and a “Start All” button to execute both analyses sequentially. During processing, a progress bar provides real-time feedback.

#### (4) Download Results

Upon completion, the “Download Options” panel allows the user to save the results. The software generates a single comma-separated values (CSV) file containing all measurements, as well as processed images and figures for each analysis, ready for import into statistical software such as R or Python.

### 2.3. Analysis Module 1: Rapid Length Estimation (OBB Method)

The “Rapid Scan” module is designed for quick, high-throughput linear length estimation across a wide range of insect taxa. This module employs a deep learning model based on the YOLOv8-obb architecture (Jocher et al., 2022/Jocher et al., 2023). We specifically chose an Oriented Bounding Box (OBB) approach because, unlike standard axis-aligned bounding boxes, an OBB can rotate to capture a specimen’s orientation. This is crucial for accurately deriving body length from the box’s longest side, regardless of how the insect is positioned in the image. To achieve high accuracy, we performed transfer learning, fine-tuning a pre-trained model on our custom-annotated dataset of 815 images from diverse insect orders (Diptera, Hymenoptera, Coleoptera). This process adapted the model to detect an OBB that tightly encloses the main body of an insect (Figure 2). Further details on model training and evaluation are available in Appendix S1.

**Figure 2.**
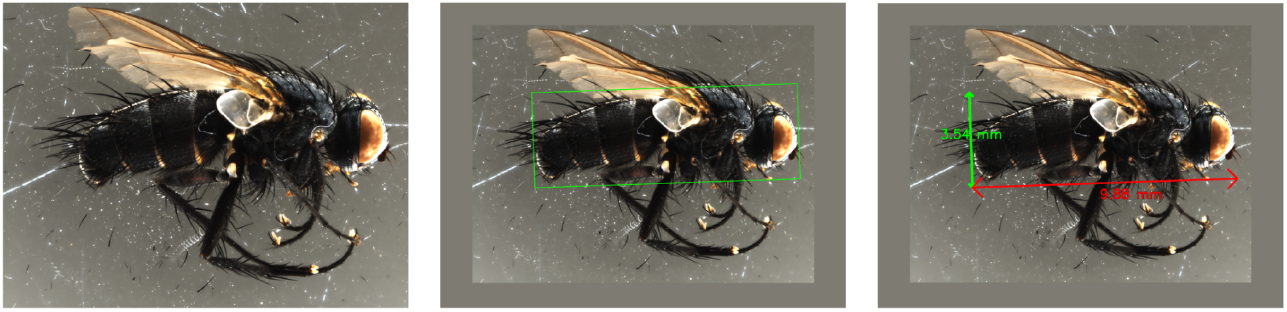
The workflow of the Rapid Length Estimation module. (left) Input image. (center) The predicted oriented bounding box (OBB). To ensure accurate OBB placement even for specimens near the image edge, a 10% padded border was added during the model training process. (right) The final output shows the derived linear length (red) and width (green) measurements.

### 2.4. Analysis Module 2: Detailed Morphometrics (Segmentation Method)

The “Detailed Analysis” module provides high-precision morphometric data for specific taxa. As an example for developing and integrating taxon-specific models, we have currently implemented a model for bristle flies (Diptera: Tachinidae), which serves to illustrate the software’s modular design. To support this extensibility, detailed guidance on the image annotation and model training workflow is provided in the software’s documentation in the GitLab repository.

The implemented module utilizes a YOLOv8-seg model, which we fine-tuned on our custom dataset of 1,320 annotated tachinid images to perform instance segmentation, precisely delineating the boundaries of the head, thorax, and abdomen (Figure 3). From these segmentation masks, the software automatically calculates the curvilinear body length by fitting a spline through the center of gravity of each body part. It then estimates the volume of each part by modeling the body as a series of stacked frustums (Wührl et al., 2022), a method that approximates 3D volume from 2D cross-sections. This module includes robust error handling; if an image of a non-target species is processed, the software will skip the image, flag it with an error message in the final CSV output, and continue with the rest of the batch. The complete model development process and the mathematical basis for the volume calculation are detailed in Appendix S1.

**Figure 3.**
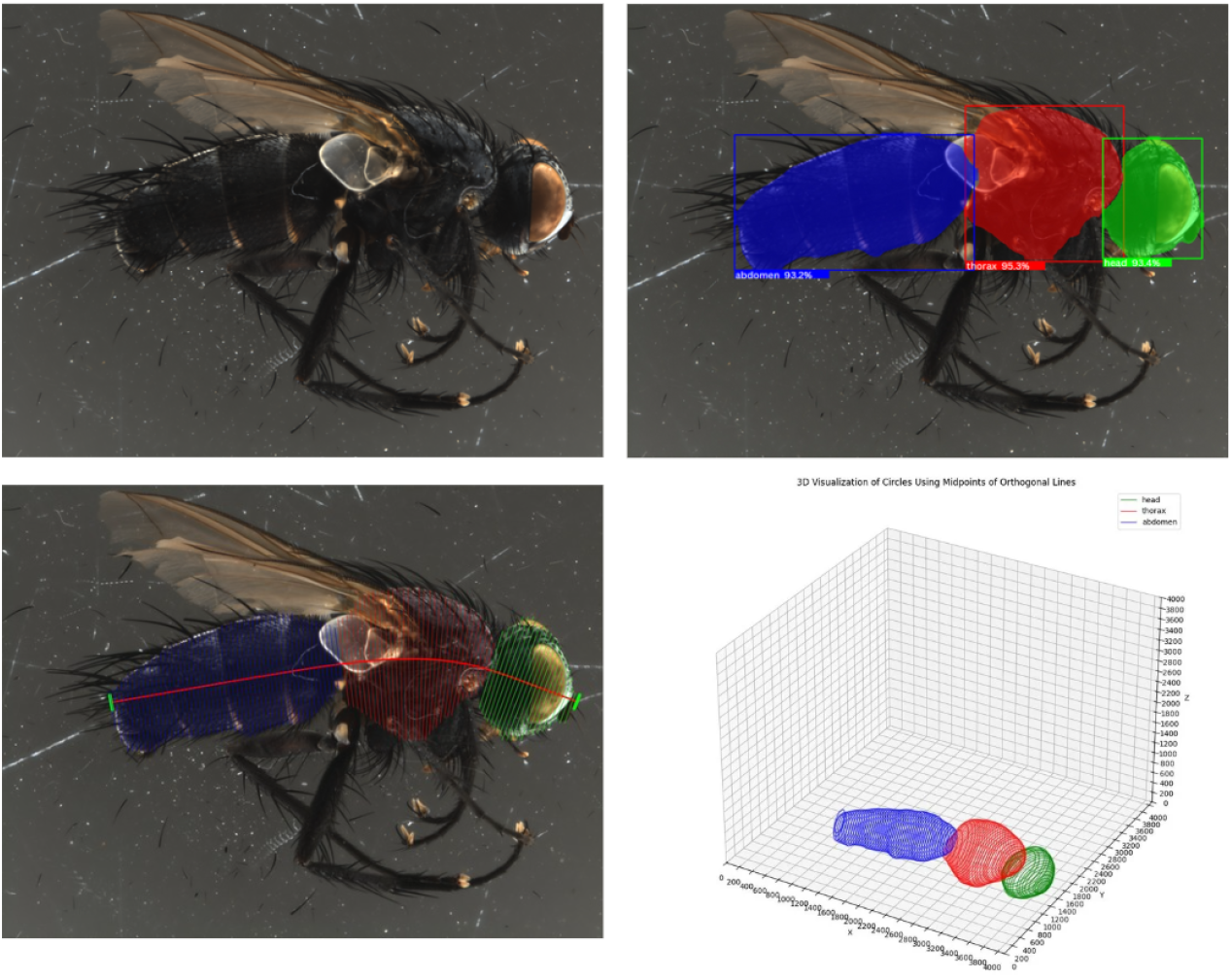
The workflow of the Detailed Morphometrics module. (top left) Input image. (top right) Predicted segmentation masks. (bottom left) Extracted the body centerline (spline) and the orthogonal cross-section lines. (bottom right) A 3D visualization of the reconstructed body volume.

## 3. Demonstration of Application

### 3.1. Validation Setup

To demonstrate the performance and accuracy of InsectMorphoAI, we conducted a validation experiment using a set of 100 tachinid fly specimens from 23 species (average length: 9.36 mm; average dry weight: 6.1 mg). For each specimen, we first captured a high-quality image using the Entomoscope setup for processing with our software. We then generated two sets of ground-truth data for comparison.

First, to validate length measurements, each specimen was re-imaged under a Zeiss Axio Zoom V16 photomicroscope. To ensure consistency, a single observer performed all manual measurements using the Zeiss software. Linear length was defined as a straight line from the anterior-most point of the head to the posterior-most point of the abdomen. Curvilinear length was measured by tracing a spline curve along the body’s central axis between the same two points. Second, to validate the volume estimates as a reliable proxy for biomass, the wet and body-only dry weights of each specimen were measured. To ensure consistency with the body-only volume estimations, the legs of each specimen were removed prior to weighing.

### 3.2. Performance and Accuracy

The “Rapid Scan” module’s linear length estimates had strong agreement with the manual Zeiss measurements, with a Mean Absolute Error (MAE) of 0.211 mm and a strong Pearson correlation coefficient (R = 0.988). To provide ecological context, this MAE represents an error of approximately 2.26% relative to the average specimen length in our validation set. Similarly, the “Detailed Analysis” module’s curvilinear length estimates were highly accurate, with an MAE of 0.309 mm (R = 0.976).

The software-derived volume exhibited a linear correlation (Figure 4) with both measured wet weight (R = 0.938, R^2^ = 0.880) and body-only dry weight (R = 0.907, R^2^ = 0.823). The software-derived volume explains over 82% of the variance in the body-only dry weight of the specimens. Using the volume in a simple linear regression model, InsectMorphoAI can predict body-only dry weight with an MAE of 0.0010 g (approximately 16.51% of the average measured dry weight).

**Figure 4.**
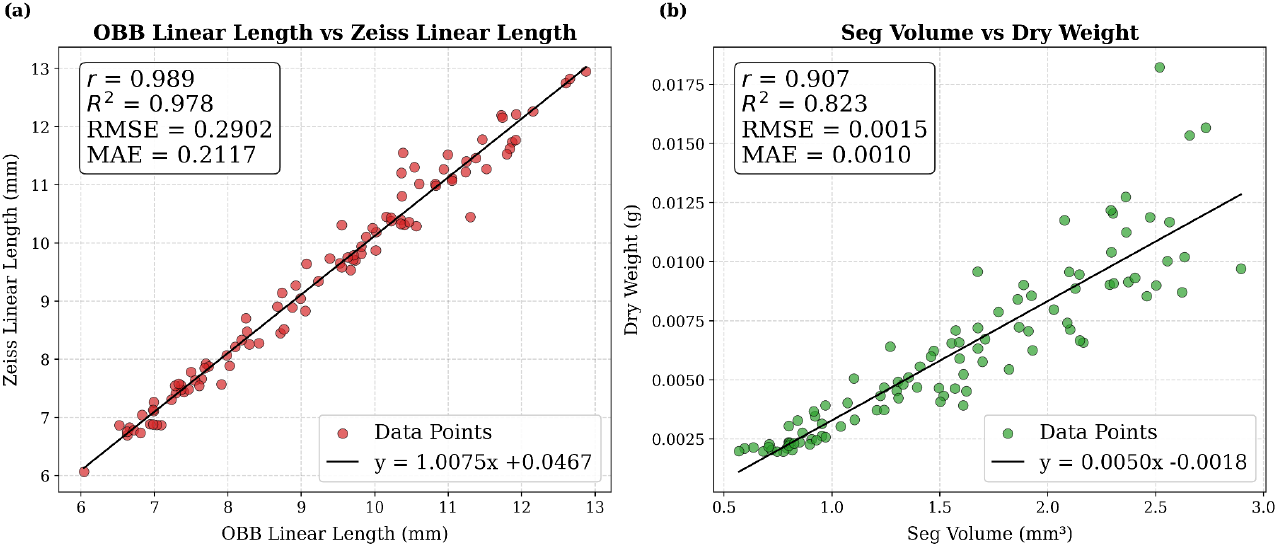
This figure demonstrates that InsectMorphoAI provides accurate length measurements and a reliable proxy for biomass. (a) A scatterplot comparing the linear length estimated by the “Rapid Scan” module versus manual Zeiss measurements. (b) A scatterplot comparing the volume estimated by the “Detailed Analysis” module against measured body-only dry weight. Both plots include the linear regression line, R^2^ value, and Mean Absolute Error (MAE).

## 4. Discussion

InsectMorphoAI provides an effective, accessible, and non-invasive software solution for a key bottleneck in ecological data collection. Automating the estimation of insect length, volume, and biomass enables researchers to gather crucial morphometric data at scales that are unfeasible with traditional manual methods. The software’s user-friendly interface and straightforward installation via Docker lower the barrier to entry for all biologists. For power users and large-scale projects, the native Python installation, along with a fully featured command-line interface, provides a path for high-performance, automated batch processing. This headless version is optimized for efficiency, leveraging multiprocessing to rapidly analyze thousands of images in a single command.

The dual-mode design is a key strength, offering a flexible toolkit for a range of research questions. The “Rapid Scan” module provides a fast and robust method for general length estimation across diverse taxa. This functionality is ideal for initial size-sorting of bulk samples from Malaise traps, for tracking broad changes in community size structure over time, or for any study where a quick size proxy is needed for thousands of specimens. In contrast, the “Detailed Analysis” module serves as a powerful tool for in-depth studies. By providing precise curvilinear length and volume estimates for specific body parts, it enables detailed research into functional trait ecology, intraspecific variation, or the population-level responses of key pest or beneficial species.

It is important to acknowledge the current limitations of the software, which also highlight avenues for future development. The high-precision segmentation model is currently specialized for the Tachinidae family. However, the software’s modular architecture is designed to support this challenge; users can train models on new families and integrate them into the application, and the GitLab repository provides documentation to guide this process. This functionality is valuable given the typical rank-abundance distributions in insect communities, where developing specialized models for the ~10 most abundant families could enable detailed analyses for approximately 50% of all captured individuals (Srivathsan et al., 2023). Furthermore, tailored models could be trained to assess other pollination-relevant traits, such as proboscis length. Our volume estimation method also provides a practical tool for studying arthropod growth, particularly in larval stages where traditional approaches are hindered by small size and rapid desiccation. Photographing live larvae, temporarily immobilized via brief cold exposure, can enable volume tracking across molting events. Our future work will focus on training and integrating these models. Currently, the volume estimation algorithm assumes a circular cross-section and excludes appendages, which underestimates total biomass. Future versions could incorporate more complex geometric models. Finally, the current models were trained exclusively on ethanol-preserved material; future work should focus on formally validating and fine-tuning the models for these different preparation types.

In conclusion, InsectMorphoAI is a valuable addition to the growing ecosystem of tools for automated biodiversity analysis. By providing a validated, open-source, and easy-to-use application, it empowers researchers to integrate detailed morphometric data into their workflows, paving the way for deeper insights into insect ecology and evolution in an era of rapid environmental change.

## Acknowledgments

Our work was supported by funding from: the Center for Integrative Biodiversity Discovery at the Museum für Naturkunde Berlin and by grant #ZF4717901SK9 of the program Natural, Artificial and Cognitive Information Processing (NACIP) of the Helmholtz-Association, Germany; Helmholtz Association Initiative and Networking Fund on the HAICORE@KIT partition; European Union–NextGenerationEU as part of the National Biodiversity Future Center, Italian National Recovery and Resilience Plan (NRRP) Mission 4 Component 2 Investment 1.4 (CUP: B83C22002950007). During the preparation of this work, the authors used AI tools to improve clarity and fluency in the writing process. After using these tools, the authors reviewed and edited the content as needed and take full responsibility for the content of the published article.

## Data Availability

The image data used for training and validating the models in this study are available on Zenodo. The InsectMorphoAI software, including source code and installation instructions, is permanently archived and publicly available on the GitLab InsectMorphoAI Repository and Zenodo. All specimen details and ground-truth validation measurements are available in the Supplementary Data file.

## Supporting Information

### Appendix S1: Model Development and Post-Processing Methods

This appendix details the technical implementation of the deep learning models and post-processing algorithms that power InsectMorphoAI.

#### Annotation and Dataset Preparation

All annotations were performed by a single expert annotator to ensure consistency across the dataset. For the taxon-specific model, a two-stage strategy was employed: an initial model was used to generate pseudo-labels on a new set of images, which were then manually corrected and refined by the expert. This process efficiently expanded the final high-quality dataset to 1,320 images for the Tachinidae model. For a complete walkthrough of our recommended annotation protocol and instructions for preparing a dataset for a new taxon, users are referred to the TRAINING_GUIDE.md file in the GitLab repository.

#### Model Training and Evaluation

All models were implemented in Python using the PyTorch framework and trained on the HAICORE high-performance computing infrastructure at the Karlsruhe Institute of

Technology (KIT), using GPU nodes equipped with NVIDIA A100 GPUs. We employed a transfer learning approach for both modules. The general length estimation model is a YOLOv8m-obb architecture fine-tuned from a model pre-trained on the DOTA dataset. The taxon-specific model is a YOLOv8m-seg architecture fine-tuned from a model pre-trained on the COCO dataset.

The selection of the medium (‘m’) model variants was based on a systematic evaluation of all five available model sizes, which determined that the ‘m’ variant offered the optimal trade-off between accuracy and computational requirements for this task (see Table S1). Key training hyperparameters included a batch size of 16 and the AdamW optimizer with a learning rate of 0.01. To ensure robustness against variations in specimen orientation and pose, we applied a comprehensive suite of data augmentations during training, including random rotations, scaling, horizontal flipping, and mosaic augmentation. The training process for both models demonstrated effective learning and generalization, as evidenced by converging precision metrics and declining loss curves (Figures S1 and S2).

#### Curvilinear Length and Volume Calculation

The centerline of the insect body is determined by calculating the centers of gravity (CoGs) of the segmented head, thorax, and abdomen. A parametric cubic spline is interpolated through these three points to create a smooth curve representing the body’s central axis. The total curvilinear length is calculated by summing the lengths of the spline segments. Volume is calculated by generating a series of orthogonal lines at equidistant intervals along the body centerline. The body is modeled as a series of frustums (truncated cones) between adjacent cross-sections. The volume *V*_*i*_ of each frustum is calculated using the formula:

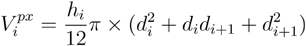

Where *h*_*i*_ is the distance between cross-sections and *d*_*i*_ and *d*_*i*+1_ are their respective diameters measured in pixels. The total volume, initially calculated in cubic pixels, is then converted to cubic millimeters using the user-defined calibration factor.

**Table S1.**
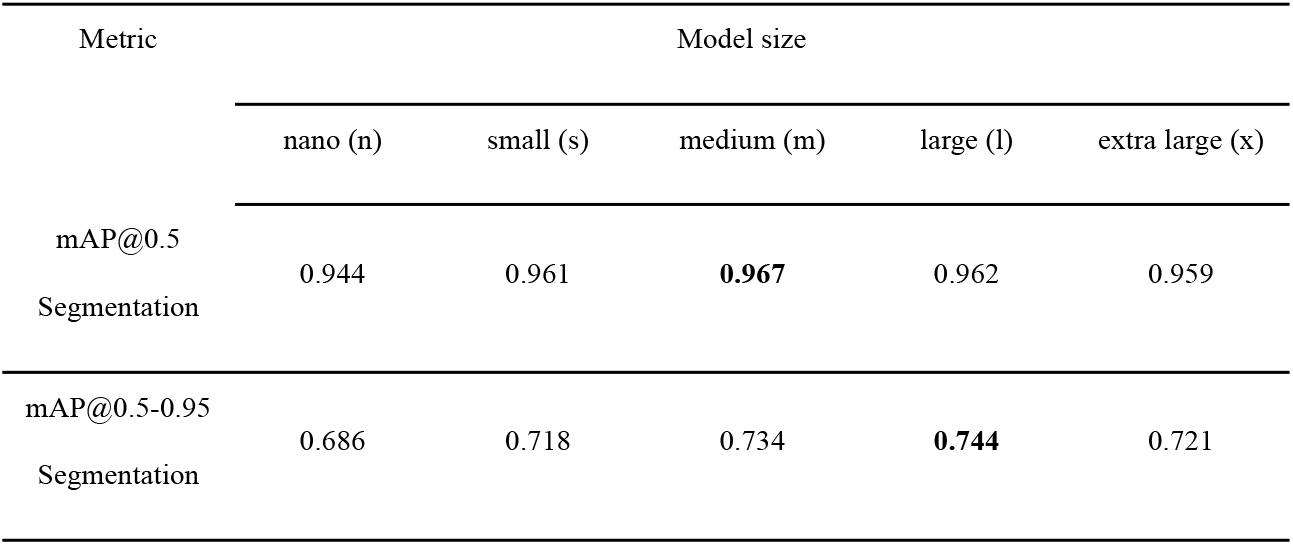
Performance comparison for each YOLOv8-seg model variant, evaluated on the initial dataset. The large (l) and medium (m) models provided the best balance between accuracy and computational requirements, informing our final model selection. mAP is the mean Average Precision.

| Metric | Model size |  |  |  |  |
| --- | --- | --- | --- | --- | --- |
|  | nano (n) | small (s) | medium (m) | large (l) | extra large (x) |
| mAP@0.5<br>Segmentation | 0.944 | 0.961 | **0.967** | 0.962 | 0.959 |
| mAP@0.5-0.95<br>Segmentation | 0.686 | 0.718 | 0.734 | **0.744** | 0.721 |

**Figure S1.**
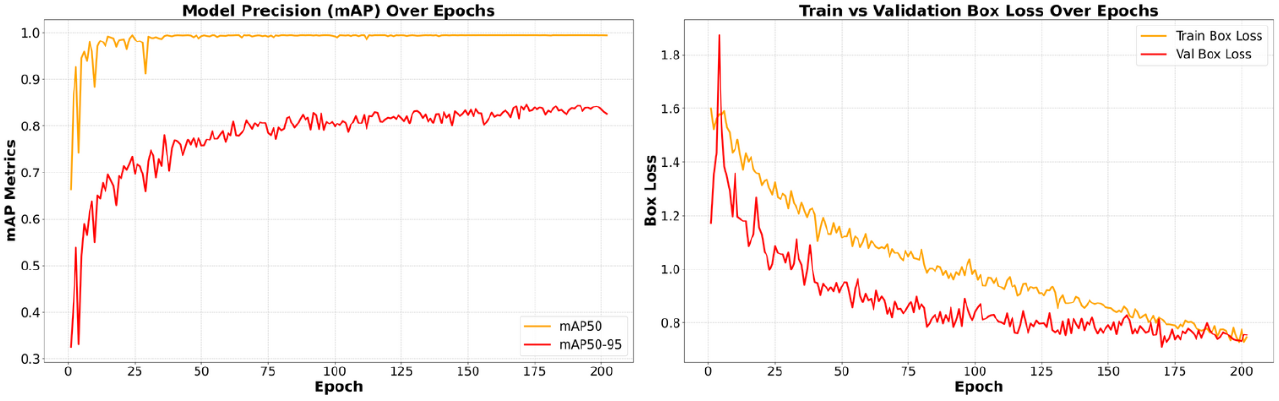
Training performance of the YOLOv8m-obb model, showing (a) model precision (mAP) and (b) box loss over 200 epochs. The convergence of the curves indicates effective learning for oriented bounding box detection.

**Figure S2.**
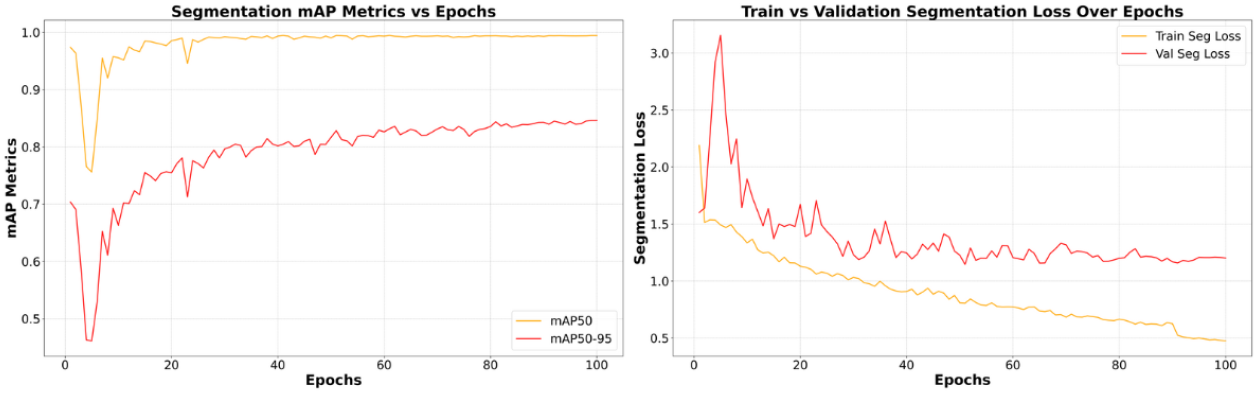
Training performance of the final YOLOv8m-seg model, showing (a) segmentation mAP and (b) segmentation loss over 100 epochs. The steady improvement in metrics and declining loss demonstrates robust training for body part segmentation.

**Figure S3.**
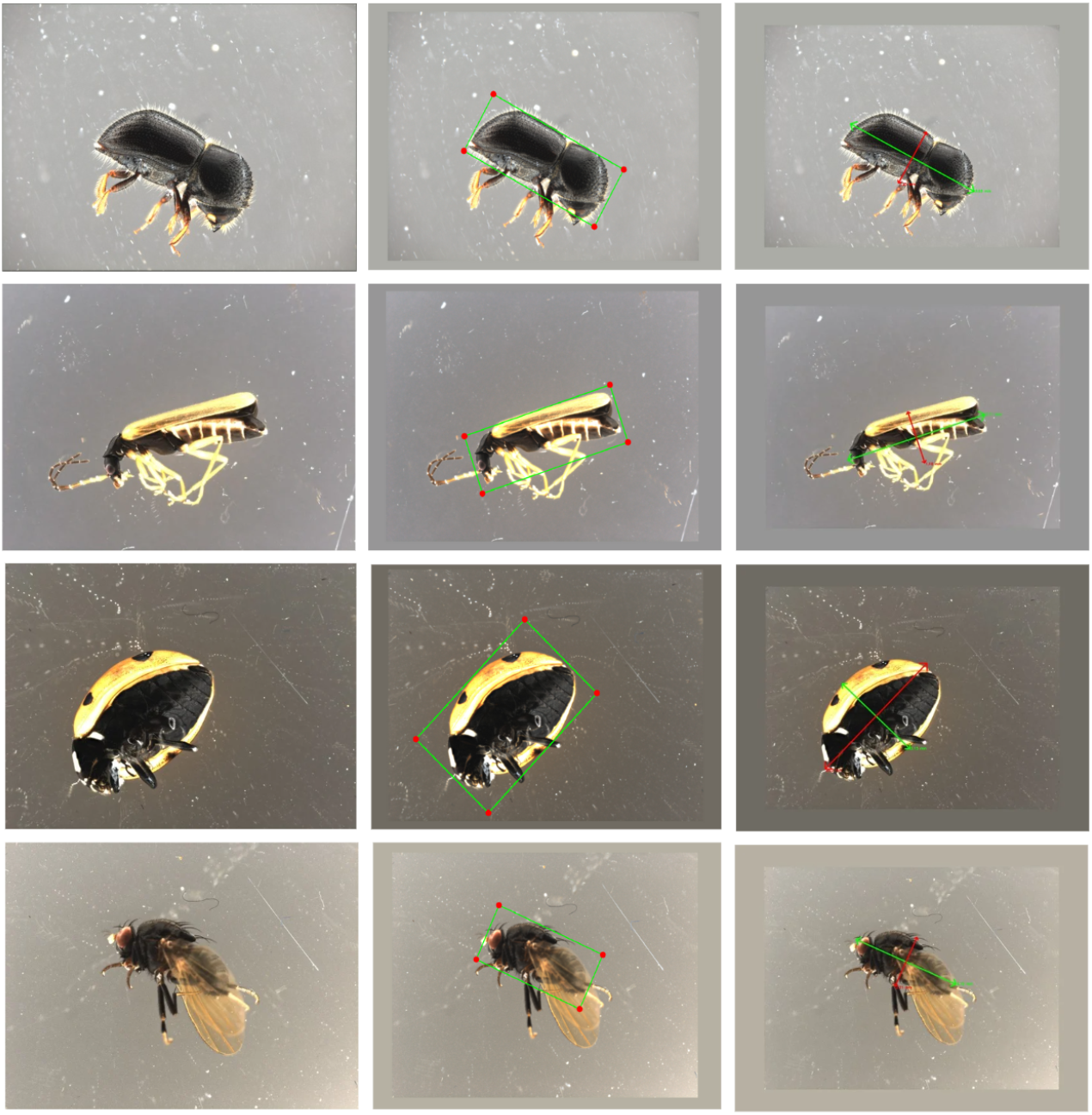
Demonstration of the general applicability of the OBB-based “Rapid Scan” module across different insect orders.

**Figure S4.**
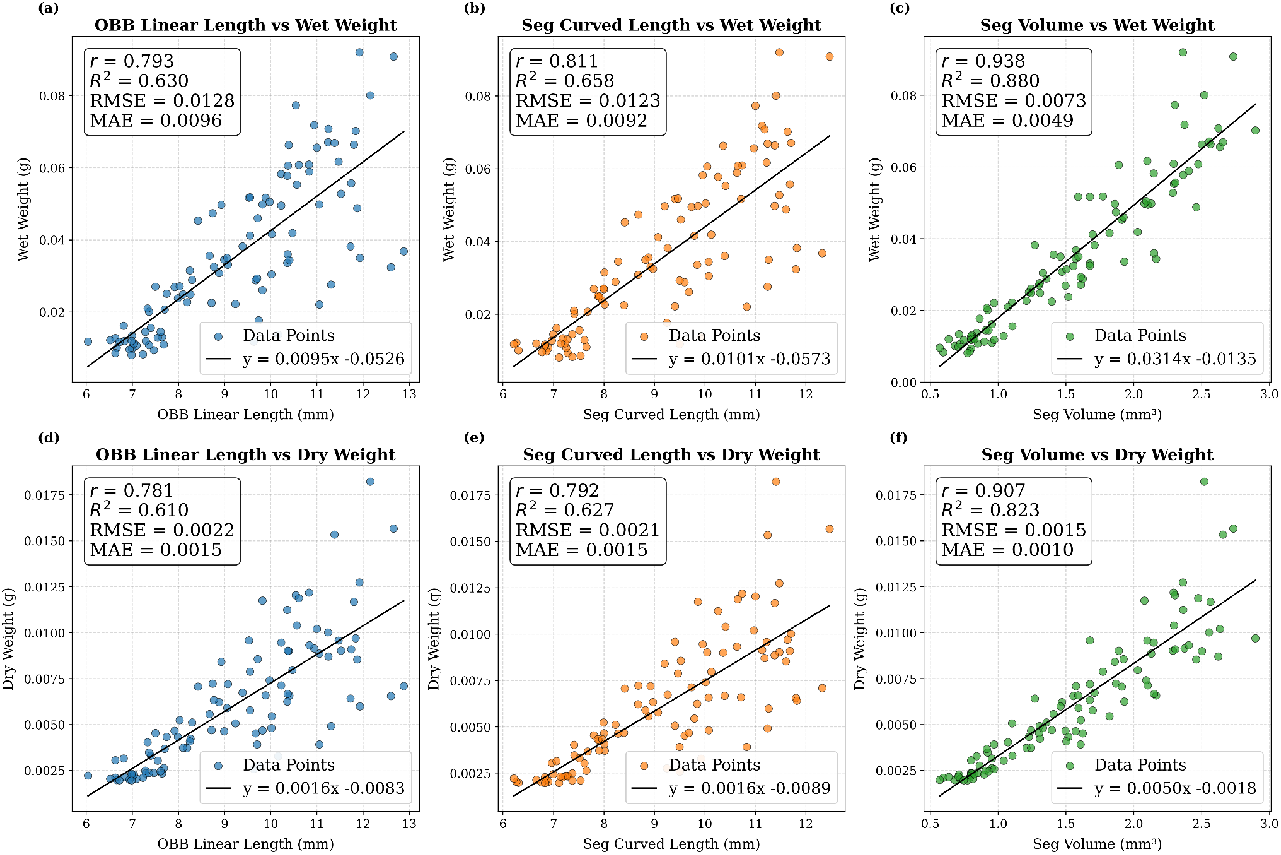
Comprehensive comparison of the linear relationships between three image-derived features (OBB linear length, Segmentation curved length, and Segmentation volume) and measured wet weight (panels a-c) and body-only dry weight (panels d-f) for 100 tachinid specimens. Each panel displays the data points, the linear regression fit, and key performance metrics (R, R^2^, RMSE, MAE).

## References

Ärje, J., Melvad, C., Jeppesen, M. R., Madsen, S. A., Raitoharju, J., Rasmussen, M. S., Iosifidis, A., Tirronen, V., Gabbouj, M., Meissner, K., & Høye, T. T. (2020). Automatic image-based identification and biomass estimation of invertebrates. Methods in Ecology and Evolution, 11(8), 922–931. 10.1111/2041-210X.13428

Ascenzi, A., Wührl, L., Feng, V., Klug, N., Pylatiuk, C., Cerretti, P., & Meier, R. (2025). EntoSieve: Automated Size-Sorting of Insect Bulk Samples to Aid Accurate Megabarcoding and Metabarcoding. Molecular Ecology Resources, n/a(n/a), e14097. 10.1111/1755-0998.14097

Bartomeus, I., Gravel, D., Tylianakis, J. M., Aizen, M. A., Dickie, I. A., & Bernard-Verdier, M. (2016). A common framework for identifying linkage rules across different types of interactions. Functional Ecology, 30(12), 1894–1903. 10.1111/1365-2435.12666

Brose, U. (2010). Body-mass constraints on foraging behaviour determine population and food-web dynamics. Functional Ecology, 24(1), 28–34. 10.1111/j.1365-2435.2009.01618.x

Cariveau, D. P., Nayak, G. K., Bartomeus, I., Zientek, J., Ascher, J. S., Gibbs, J., & Winfree, R. (2016). The Allometry of Bee Proboscis Length and Its Uses in Ecology. PLOS ONE, 11(3), e0151482. 10.1371/journal.pone.0151482

Caruso, V., Shirali, H., Bouget, C., Cerretti, P., Curletti, G., Groot, M. de, Groznik, E., Gutowski, J. M., Pyliatuk, C., Roques, A., Sallé, A., Sweeney, J., Wührl, L., & Rassati, D. (2025). Image-based recognition using advanced neural networks can aid surveillance of Agrilus (Coleoptera, Buprestidae) jewel beetles. ARPHA Preprints, 6, e154842. 10.3897/arphapreprints.e154842

Fox, C. W., & Czesak, M. E. (2000). Evolutionary Ecology of Progeny Size in Arthropods. Annual Review of Entomology, 45(Volume 45, 2000), 341–369. 10.1146/annurev.ento.45.1.341

Greenleaf, S. S., Williams, N. M., Winfree, R., & Kremen, C. (2007). Bee foraging ranges and their relationship to body size. Oecologia, 153(3), 589–596. 10.1007/s00442-007-0752-9

Hallmann, C. A., Sorg, M., Jongejans, E., Siepel, H., Hofland, N., Schwan, H., Stenmans, W., Müller, A., Sumser, H., Hörren, T., Goulson, D., & Kroon, H. de. (2017). More than 75 percent decline over 27 years in total flying insect biomass in protected areas. PLOS ONE, 12(10), e0185809. 10.1371/journal.pone.0185809

Hartop, E., Lee, L., Srivathsan, A., Jones, M., Peña-Aguilera, P., Ovaskainen, O., Roslin, T., & Meier, R. (2024). Resolving biology’s dark matter: Species richness, spatiotemporal distribution, and community composition of a dark taxon. BMC Biology, 22(1), 215. 10.1186/s12915-024-02010-z

Hereld, M., Ferrier, N. J., Agarwal, N., & Sierwald, P. (2017). Designing a High-Throughput Pipeline for Digitizing Pinned Insects. 2017 IEEE 13th International Conference on E-Science (e-Science), 542–550. 10.1109/eScience.2017.88

Jocher, G., Chaurasia, A., & Qiu, J. (2023). Ultralytics YOLO (Version 8.0.0) [Python]. https://github.com/ultralytics/ultralytics (Original work published 2022)

Karlsson, D., Forshage, M., Holston, K., & Ronquist, F. (2020). The data of the Swedish Malaise Trap Project, a countrywide inventory of Sweden’s insect fauna. Biodiversity Data Journal, 8, e56286. 10.3897/BDJ.8.e56286

Kendall, L. K., Rader, R., Gagic, V., Cariveau, D. P., Albrecht, M., Baldock, K. C. R., Freitas, B. M., Hall, M., Holzschuh, A., Molina, F. P., Morten, J. M., Pereira, J. S., Portman, Z. M., Roberts, S. P. M., Rodriguez, J., Russo, L., Sutter, L., Vereecken, N. J., & Bartomeus, I. (2019). Pollinator size and its consequences: Robust estimates of body size in pollinating insects. Ecology and Evolution, 9(4), 1702–1714. 10.1002/ece3.4835

Klug, N., Kramer, M., Mazrek, F., Wührl, L., Shirali, H., Meier, R., & Pylatiuk, C. (2024). Automated Photogrammetric Close-Range Imaging System for Small Invertebrates Using Acoustic Levitation. 10.36227/techrxiv.172651022.21831566/v1

Lister, B. C., & Garcia, A. (2018). Climate-driven declines in arthropod abundance restructure a rainforest food web. Proceedings of the National Academy of Sciences, 115(44), E10397–E10406. 10.1073/pnas.1722477115

Meier, R., Lawniczak, M. K. N., & Srivathsan, A. (2025). Illuminating Entomological Dark Matter with DNA Barcodes in an Era of Insect Decline, Deep Learning, and Genomics. Annual Review of Entomology, 70(Volume 70, 2025), 185–204. 10.1146/annurev-ento-040124-014001

Miller-ter Kuile, A., Apigo, A., Bui, A., DiFiore, B., Forbes, E. S., Lee, M., Orr, D., Preston, D. L., Behm, R., Bogar, T., Childress, J., Dirzo, R., Klope, M., Lafferty, K. D., McLaughlin, J., Morse, M., Motta, C., Park, K., Plummer, K., … Young, H. (2022). Predator–prey interactions of terrestrial invertebrates are determined by predator body size and species identity. Ecology, 103(5), e3634. 10.1002/ecy.3634

Sánchez-Bayo, F., & Wyckhuys, K. A. G. (2019). Worldwide decline of the entomofauna: A review of its drivers. Biological Conservation, 232, 8–27. 10.1016/j.biocon.2019.01.020

Schneider, S., Taylor, G. W., Kremer, S. C., Burgess, P., McGroarty, J., Mitsui, K., Zhuang, A., deWaard, J. R., & Fryxell, J. M. (2022). Bulk arthropod abundance, biomass and diversity estimation using deep learning for computer vision. Methods in Ecology and Evolution, 13(2), 346–357. 10.1111/2041-210X.13769

Shirali, H., Hübner, J., Both, R., Raupach, M., Reischl, M., Schmidt, S., & Pylatiuk, C. (2024). Image-based recognition of parasitoid wasps using advanced neural networks. Invertebrate Systematics, 38(6). 10.1071/IS24011

Srivathsan, A., Ang, Y., Heraty, J. M., Hwang, W. S., Jusoh, W. F. A., Kutty, S. N., Puniamoorthy, J., Yeo, D., Roslin, T., & Meier, R. (2023). Convergence of dominance and neglect in flying insect diversity. Nature Ecology & Evolution, 7(7), Article 7. 10.1038/s41559-023-02066-0

Srivathsan, A., Lee, L., Katoh, K., Hartop, E., Kutty, S. N., Wong, J., Yeo, D., & Meier, R. (2021). ONTbarcoder and MinION barcodes aid biodiversity discovery and identification by everyone, for everyone. BMC Biology, 19(1), 217. 10.1186/s12915-021-01141-x

Streamlit Inc. (n.d.). Streamlit: A faster way to build and share data apps [Computer software]. Retrieved September 3, 2025, from https://streamlit.io/

Wägele, J. W., Bodesheim, P., Bourlat, S. J., Denzler, J., Diepenbroek, M., Fonseca, V., Frommolt, K.-H., Geiger, M. F., Gemeinholzer, B., Glöckner, F. O., Haucke, T., Kirse, A., Kölpin, A., Kostadinov, I., Kühl, H. S., Kurth, F., Lasseck, M., Liedke, S., Losch, F., … Wildermann, S. (2022). Towards a multisensor station for automated biodiversity monitoring. Basic and Applied Ecology, 59, 105–138. 10.1016/j.baae.2022.01.003

Wührl, L., Pylatiuk, C., Giersch, M., Lapp, F., von Rintelen, T., Balke, M., Schmidt, S., Cerretti, P., & Meier, R. (2022). DiversityScanner: Robotic handling of small invertebrates with machine learning methods. Molecular Ecology Resources, 22(4), 1626–1638. 10.1111/1755-0998.13567

Wührl, L., Rettenberger, L., Meier, R., Hartop, E., Graf, J., & Pylatiuk, C. (2024). Entomoscope: An Open-Source Photomicroscope for Biodiversity Discovery. IEEE Access, 12, 11785–11794. IEEE Access. 10.1109/ACCESS.2024.3355272

